# Comparative Analysis of AAV Serotypes for Transduction of Olfactory Sensory Neurons

**DOI:** 10.1101/2024.09.26.615247

**Authors:** Benjamin D.W. Belfort, Johnathan D. Jia, Alexandra Garza, Anthony M. Insalaco, JP McGinnis, Brandon T. Pekarek, Joshua Ortiz-Guzman, Burak Tepe, Hu Chen, aSCENT-PD Investigators, Zhandong Liu, Benjamin R. Arenkiel

## Abstract

Olfactory sensory neurons within the nasal epithelium detect volatile odorants and relay odor information to the central nervous system. Unlike other sensory inputs, olfactory sensory neurons interface with the external environment and project their axons directly into the central nervous system. The use of adeno-associated viruses to target these neurons has garnered interest for applications in gene therapy, probing olfactory sensory neuron biology, and modeling disease. To date, there is no consensus on the optimal AAV serotype for efficient and selective transduction of olfactory sensory neurons *in vivo*. Here we utilized serial confocal imaging and single-nucleus RNA sequencing to evaluate the efficacy of 11 different AAV serotypes in transducing murine olfactory sensory neurons via non-invasive nasal inoculation. Our results reveal that AAV1, while highly effective, exhibited broad tropism, whereas AAV-DJ/8 showed the greatest specificity for olfactory sensory neurons.

## Introduction

The mammalian olfactory epithelium (OE) is a specialized epithelial tissue within the nose that plays a critical role in the sense of smell. Olfactory sensory neurons (OSNs) are specialized chemosensory neurons that reside in the OE and detect airborne chemicals through G-protein coupled odorant receptors. OSNs project their axons directly into the olfactory bulb (OB), creating a single-neuron chain between the external environment and the central nervous system (CNS). This conduit into the CNS provides unique opportunities for probing OSN biology, administering targeted gene therapies, and modeling disease. The use of AAVs has been explored for these purposes ^1^, yet information is lacking on the optimal serotype for transducing OSNs. In the present study, we leveraged confocal imaging and single-nucleus RNA sequencing (snRNAseq) to identify optimal AAV serotypes for *in vivo* transduction of murine OSNs.

## Results

To test the efficacy of AAV transduction across commonly used serotypes, we packaged an identical construct (rAAV-EF1a-TdTomato-WPRE-PolyA) into 11 AAV different capsid serotypes with reported OE and neuronal transduction^1^. Serotypes included AAV1, AAV2, AAV5, AAV7, AAV8, AAV9, AAV-DJ/8 ^2^,^3^ AAV-PhP.eB ^4^ , AAV-PhP.S ^4^ , AAV-rh10 ^5,6^ , and AAV-SCH9 ^7^. Each AAV serotype was introduced via non-invasive nasal inoculation (NINI) ^8^ into three male mice per serotype, for a total of 33 experimental animals. After a four-week expression period, the OB was processed to quantify colocalized olfactory marker protein (OMP) and TdTomato expression within OSN axon terminals as a readout for the number of transduced OSNs [Fig. 1A]. By first focusing on the reporter expression in the glomerular layer of the OB, we avoided potentially confounding TdTomato signals from other transduced accessory cell types within the OE [Fig. 1B].

**Figure 1.**
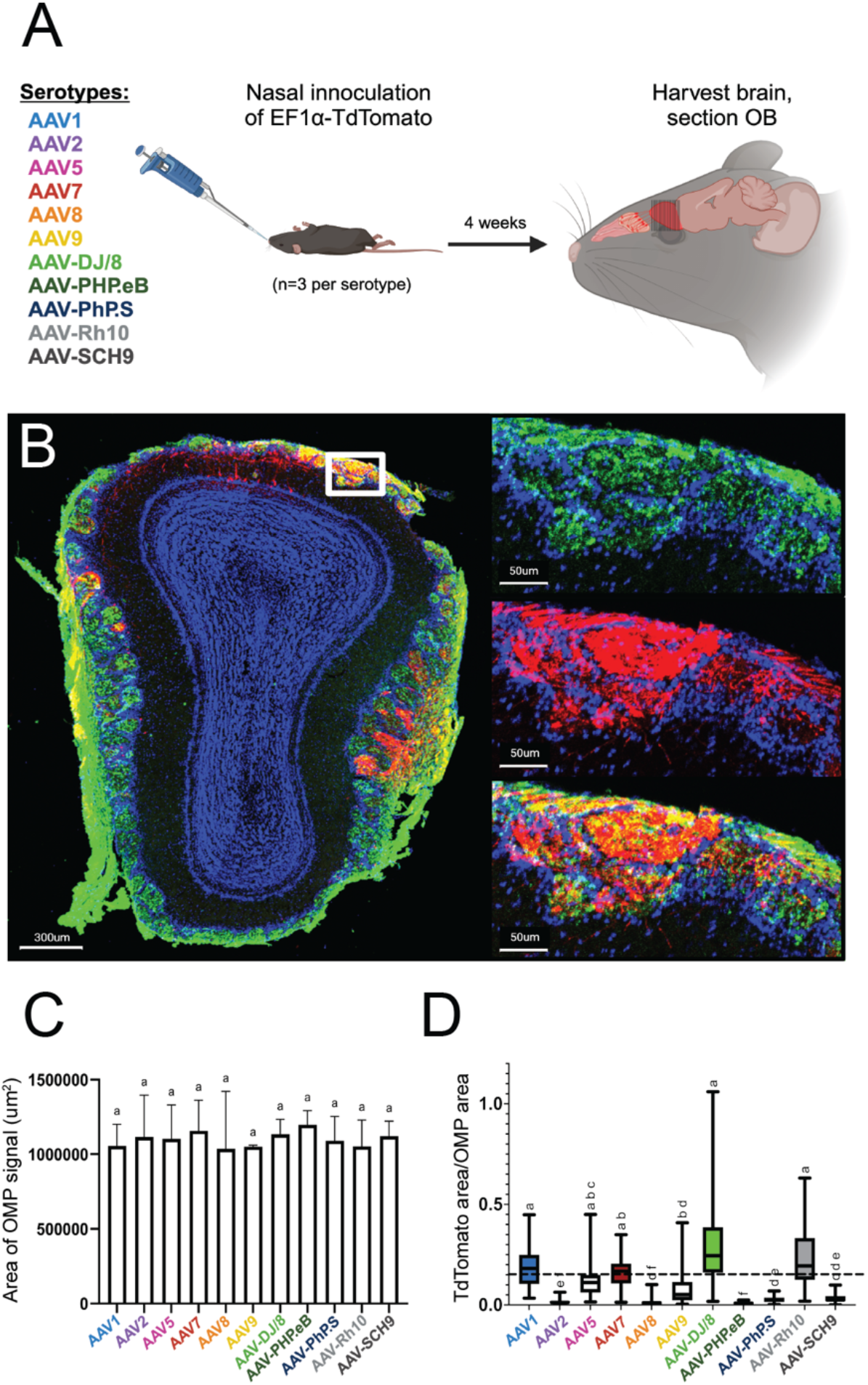
Confocal imaging identifies AAV1, AAV7, AAV-DJ/8, and AAV-rh10 as efficient AAV serotypes for OSN transduction. A) Overview of experimental paradigm. Identical plasmid constructs were packaged in 11 unique AAV serotypes, and each serotype was individually introduced via NINI. B) Representative confocal image of OB cross-section from an AAV1 inoculated mouse. Upper right: close-up of OSNs forming the structures of the glomerular and olfactory nerve layers of the OB, with OMP (green) channel isolated. Middle right: close-up of the same region, with TdTomato (red) channel isolated. Bottom right: overlay of OMP and TdTomato channels. Blue= Hoechst. C) Total area of OMP signal per OB for each serotype. D) Quantification of the percentage of TdTomato/OMP in the glomerular layer per OB section. Error bars represent the minimum and maximum values. Statistical differences were assessed using one-way ANOVA, the results of which are represented by compact letter display.

For quantification, we performed confocal image analysis across OSN terminal fields within the glomerular layer via immunostaining, measuring levels of tdTomato reporter expression alongside OMP. Importantly, we found no difference in total OMP area between samples from each serotype [Fig. 1C], suggesting that transduction itself did not compromise overall OSN viability or OE integrity. We next quantified the amount of TdTomato surface area (normalized to OMP surface area) per section, and found that AAV1, AAV7, AAV-DJ/8, and AAV-rh10 displayed the greatest levels of expression [Fig. 1D] in OSN terminals that project to the olfactory bulb.

Following this primary reporter expression screen, we next performed snRNAseq to more accurately quantify OSN transduction efficiency of the top 4 candidates in the different OE cell types. Towards this, we designed 4 new AAV expression constructs, all identical aside from a short barcode sequence (“ID”) that identifies the corresponding AAV serotype upon transduction and snRNA sequencing. The 4 AAVs were mixed at equal titer and introduced into wildtype mice via NINI. After 4 weeks of *in vivo* expression, OE was harvested for snRNAseq [Fig. 2A]. Data were processed following standard scRNAseq guidelines ^9^. Using 20,081 high-quality nuclei, we generated an annotated UMAP from known cell type-specific markers (see SI appendix) to quantify the distribution of cell types [Fig. 2B-D] and identify those that were differentially transduced by the four candidate AAV serotypes. We confirmed cell type lineages by comparing terminal fate probabilities from Palantir pseudotime analysis, with Horizontal Basal Cells (HBCs) designated as the starting point and mature OSN (mOSN), mature sustentacular (mSus), microvillar (MVC), and Bowman’s gland (BowG) cells as terminal fates ^10^ [Fig. E-F]. A 2-sided t-test (α = 0.05) and chi-squared goodness of fit indicated that the terminal fate probabilities between GBCs and iSus cells were different for mOSN and mSus lineages [Fig. 2G]. Finally, we quantified the number and type of cells that were effectively transduced for each AAV serotype [Fig. 2H]. In total, we identified 382 cells that harbored AAV barcodes. While mOSN transduction efficiency was highest for AAV1 [Fig. 2I], AAV-DJ/8 showed the greatest specificity for mOSNs [Fig. 2J]. Further examination of AAV1 expression showed broad tropism, with immature sustentacular (iSus) and airway ciliated cells (ACCs) contributing to the highest normalized cell counts [Fig. 2K].

**Figure 2.**
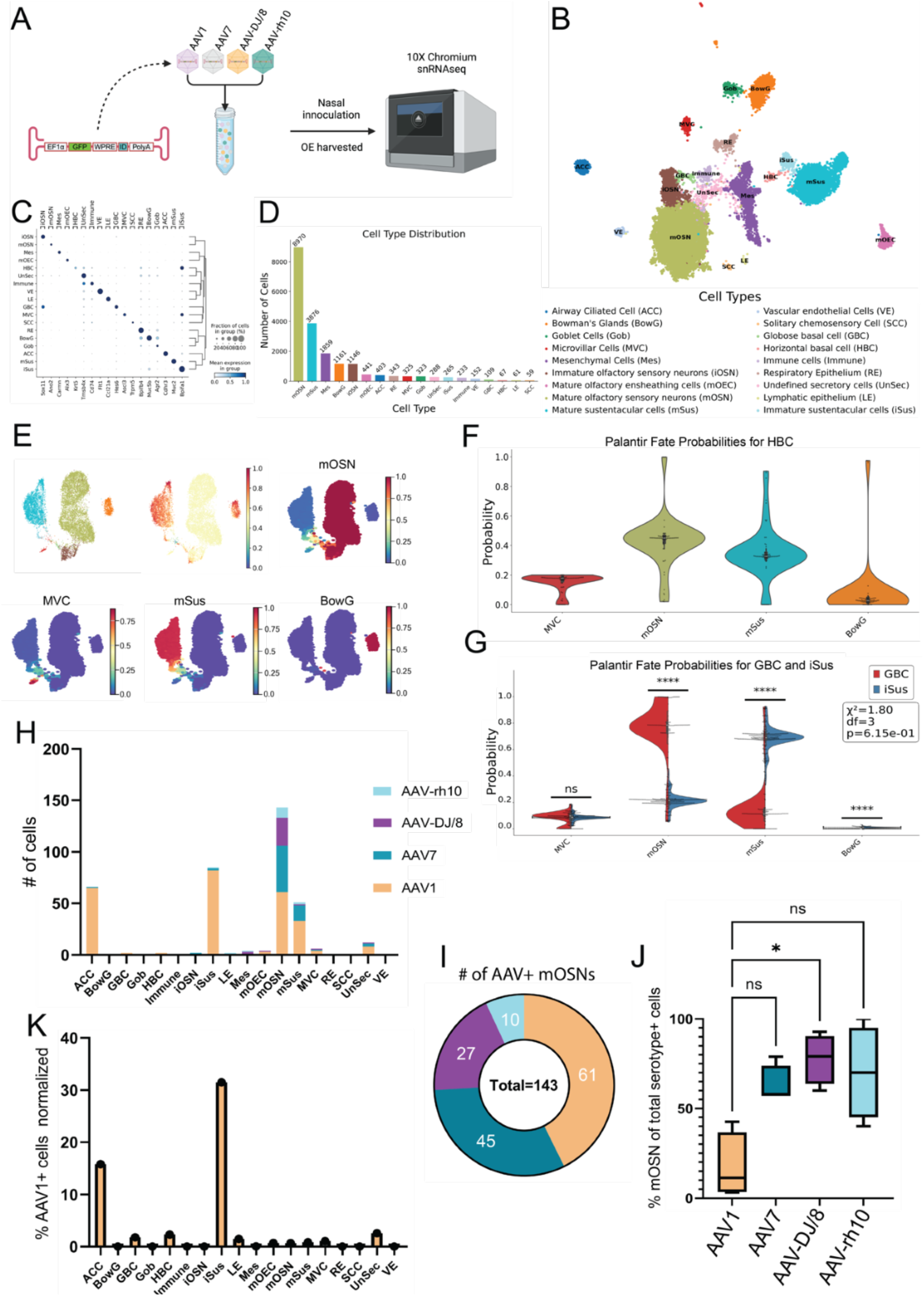
Single nucleus transcriptomics shows AAV1 has the greatest transduction rate while AAV-DJ/8 has the greatest specificity for mOSNs. A) Experimental workflow. B) UMAP of snRNAseq dataset. C) Dot plot showing the cell-type specific markers used for annotations. The x-axis is the marker, and the y-axis is the cell type. D) Cell type distribution with total cell counts. The x-axis is the cell type with each bar’s color reflecting the corresponding color on the UMAP, and the y-axis is the absolute count of the respective cell types. E) Minimum distortion embedding (MDE) with our specific lineages of interest including mOSN, mSus, MVC, and BowG lineages. The total OE cell type-specific pseudotime is shown above and individual lineage pseudotimes shown below, with red indicating late pseudotime and dark blue indicating early pseudotime. F) Violin-box plot of terminal fate probability of HBCs calculated from the Palantir pseudotime. G) Terminal fate probabilities of GBCs and iSus were compared using a 2- sided t-test (α = 0.05) and chi-squared goodness of fit for each possible terminal fate. The stars represent the level of significance. If p < 1e^-4^ there are 4 stars, < 1e^-3^ are 3 stars, 2 stars for < 0.01, 1 star for < 0.05, and ns for not significant. Chi-squared GOF was used to evaluate overall differences between terminal fate probabilities of each cluster. H) Total AAV+ cell counts per cell type. I) Total AAV+ mOSN cell counts. J) Percent of total serotype positive cells which are mOSNs. * p < .05, Kruskal-Wallis test. K) Percent AAV1+ cell counts of each cell type cluster total cell count.

## Discussion

In the present study, we leveraged confocal imaging and snRNAseq to identify optimal AAV serotypes for *in vivo* OSN transduction via NINI. Despite variability in our imaging dataset, attributable to unpredictable flow of fluid through the nasal turbinates during NINI, AAV1, AAV7, AAV-DJ/8, and AAV-rh10 displayed the greatest efficacy in their ability to transduce OSNs compared to other AAV serotypes we tested [Fig. 1D]. To determine which serotype was most efficient at transducing OSNs in a more quantitative, higher-resolution, and cell type-specific manner, we employed snRNA sequencing from isolated OE. Though this is the first study to employ snRNAseq of the OE, we found that clustering and cell type distribution were similar to other single-cell sequencing studies previously performed on the OE ^11,12, 13^ [Fig. 2B-D]. We found that cell subsets comprised known lineages of interest, namely HBCs, globose basal cells (GBCs), immature OSNs (iOSNs), mOSNs, iSus, mSus, MVCs, and BowGs, and determined the corresponding pseudotemporal lineage patterns [Fig. 2E]. Through this, we substantiated cell type identities by comparing their terminal fate probabilities. Of interest, the terminal fate probability distribution of HBCs properly reflected multiple cell types, indicating their pluripotent ability to divide into all lineages of interest, but favoring mOSNs, mSus, and MVCs [Fig. 2F]. By comparing the terminal fates of iSus and GBC, we further validated the identified OSN and sustentacular cell lineages. Having confirmed the cellular composition of the transduced tissue, we next quantified the number of AAV+ cells within each cell type cluster [Fig. 2H]. Of the serotypes tested, AAV1 showed the highest transduction efficiency of OSNs [Fig. 2I], but AAV-DJ/8, AAV7, and AAV-rh10 all showed greater OSN specificity, with AAV-DJ/8 showing the greatest specificity compared to AAV1 [Fig. 2J]. From these data, we also found that AAV1 showed the broadest tropism in the OE; transducing cell types such as HBCs, GBCs, ACCs, and particularly iSus, all with greater efficiency than OSNs [Fig. 2K]. Alongside a census of the cell types that comprise the mouse olfactory epithelium, together these findings also present useful implications for future research, including OSN-targeted gene therapy, probing OSN biology, modeling diseases where the OE and OB are loci of interest (e.g., Parkinson’s disease and Alzheimer’s disease), and informing the development of future AAV capsids for targeting OSNs.

## Materials and Methods

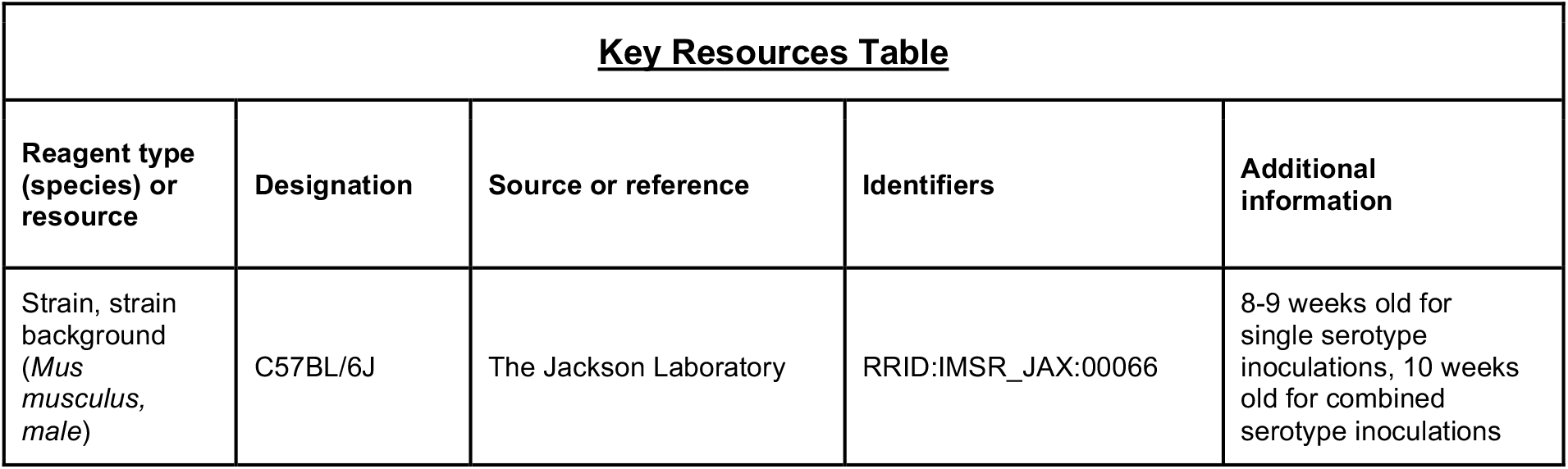
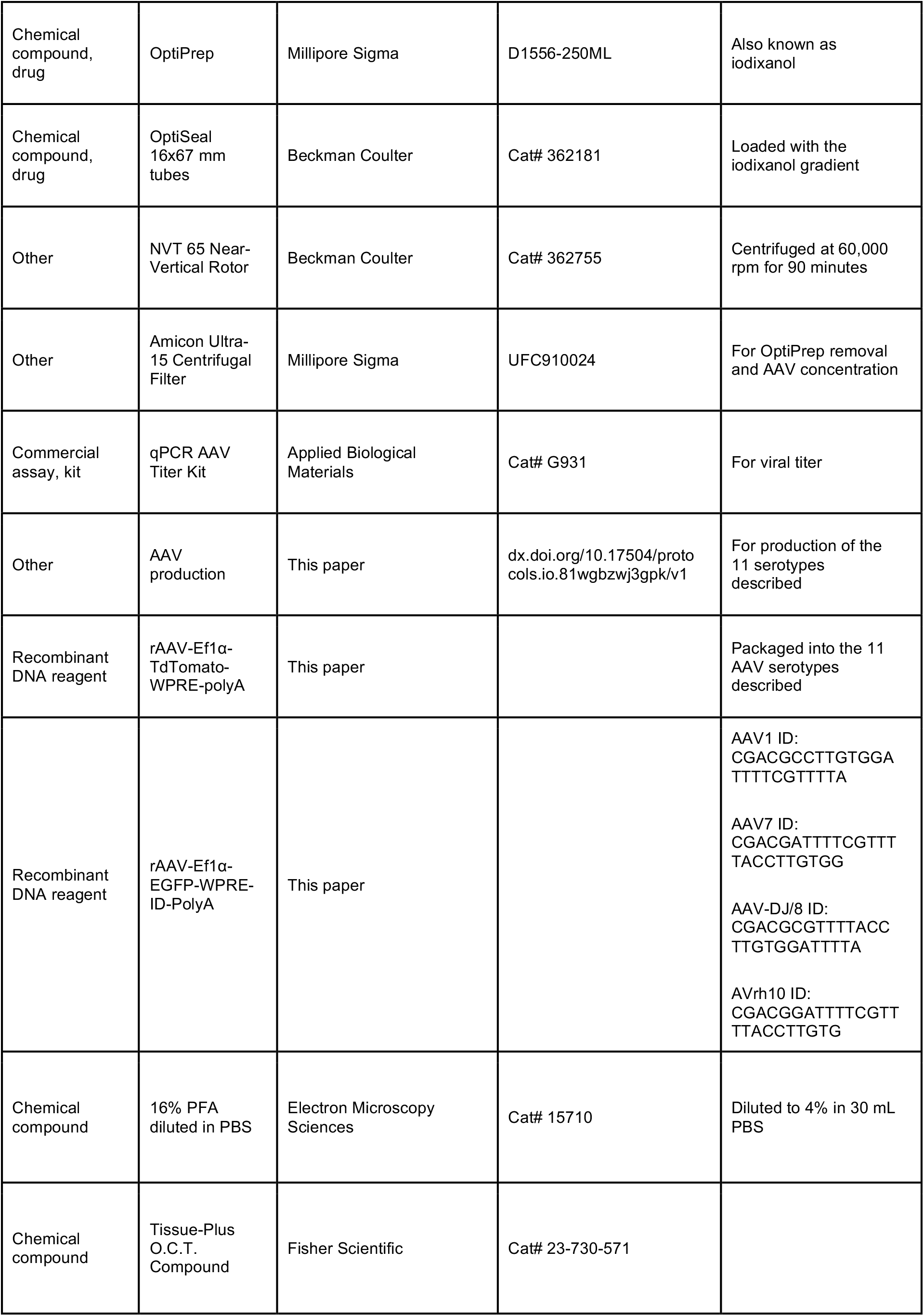
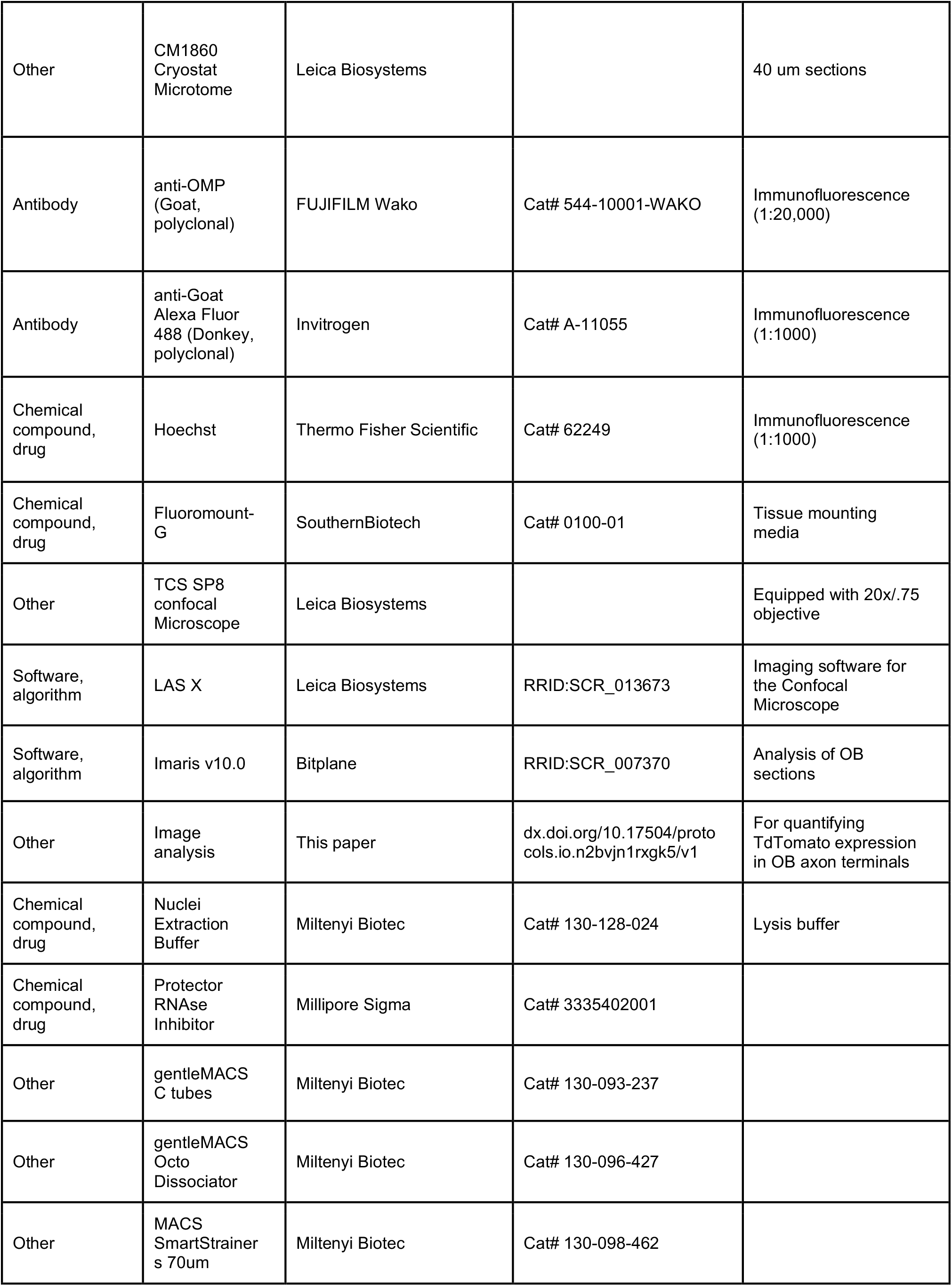
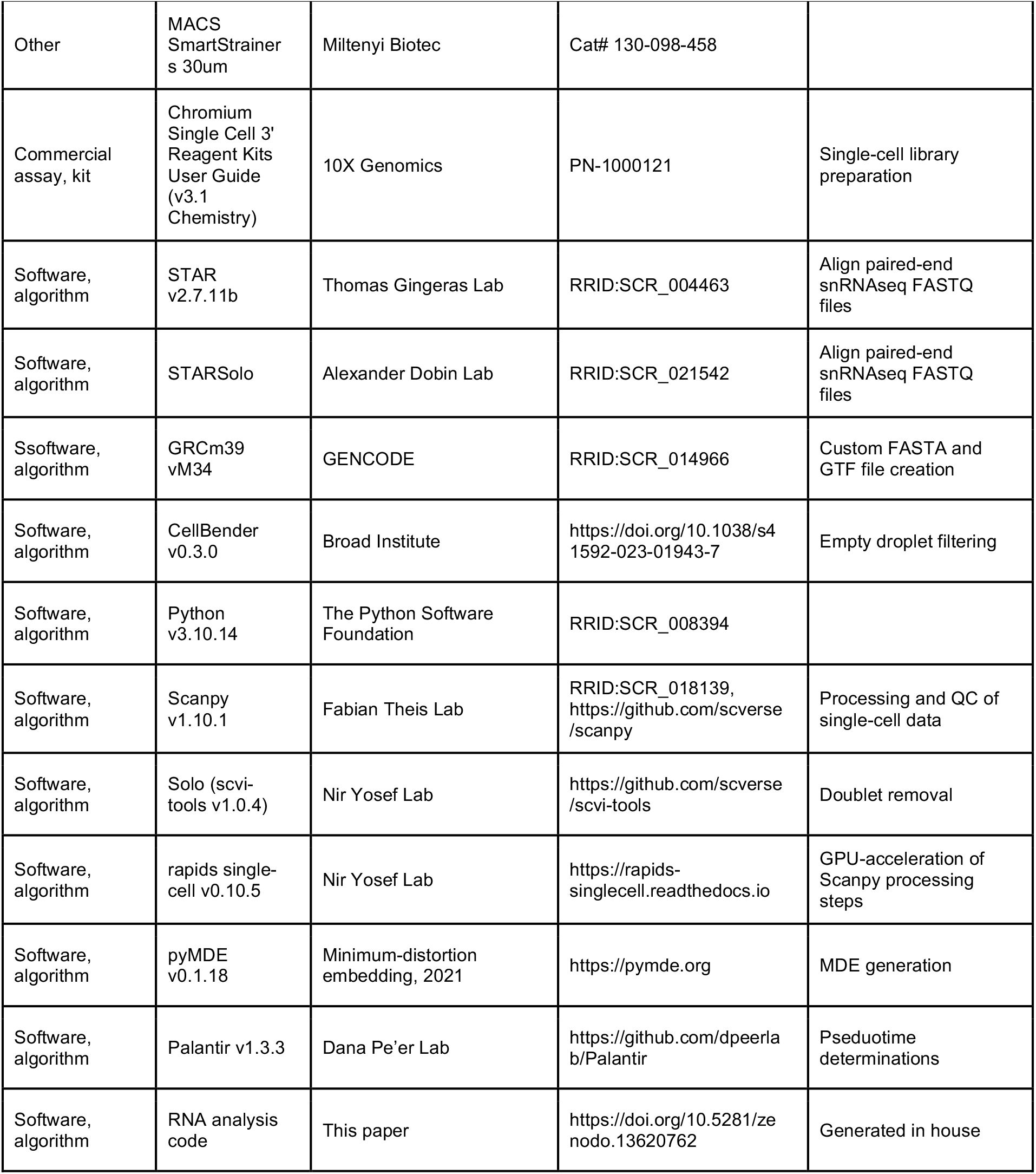

### Animals

All mice used in this study were male C57BL/6J (Jax :000664) and were used in compliance with Baylor College of Medicine IACUC. For single serotype inoculations, mice were 8 to 9 weeks old, and for combined serotype inoculations, mice were 10 weeks old.

### Plasmid Constructs and Nasal Inoculation

For all nasal inoculations, mice were briefly anesthetized with isoflurane. While under anesthesia, the mice were nasally inoculated in 5 ul doses for a total of 40 ul (alternating between nostrils, with 20 ul of virus per nostril).

For inoculations of individual serotypes, rAAV-Ef1α-TdTomato-WPRE-polyA was packaged in serotype AAV1, AAV2, AAV5, AAV7, AAV8, AAV9, AAV-DJ/8, AAV-PHP.eB, AAV-PHP.S, AAVrh10, or AAV-SCH9 and were normalized to a final concentration of 4.85e^11^ vg/mL. 3 mice were inoculated with each individual serotype, for a total of 33 mice.

For mixed inoculations, unique variants of an rAAV-Ef1α-EGFP-WPRE-ID-PolyA construct were packaged in AAV1, AAV7, AAV-DJ/8, and AAVrh10, with unique barcodes (ID) corresponding to each serotype. The AAVs were combined in equal concentrations (a total of 6.10e^9^ vg per serotype). The IDs are listed below:

AAV1 ID: CGACGCCTTGTGGATTTTCGTTTTA

AAV7 ID: CGACGATTTTCGTTTTACCTTGTGG

AAV-DJ/8 ID: CGACGCGTTTTACCTTGTGGATTTT

AAVrh10 ID: CGACGGATTTTCGTTTTACCTTGTG

### Tissue Processing for Immunofluorescence

Four weeks after nasal inoculation with individual AAV serotypes, the 33 mice were deeply anesthetized with isoflurane and transcardially perfused with 10mL of ice-cold PBS followed by 10mL of 4% paraformaldehyde (16% PFA diluted in PBS; Electron Microscopy Sciences 15710). The olfactory bulbs (OBs) were then harvested, fixed in 4% PFA at 4°C overnight, and moved to 30% sucrose (36 hours) for cryoprotection. OBs were frozen in OCT (Fisher HealthCare cat# 23-730-571) and sectioned at 40um on a cryostat (Leica CM1860), with every third section collected for analysis. Tissue sections were blocked in 5% Donkey Serum (1X PBS, 0.1% Triton) for one hour at room temperature, washed three times in PBST (0.1% Triton) and incubated with anti-OMP (1:20,000, FUJIFILM Wako Chemicals U.S.A. Corporation cat# 544-10001-WAKO) antibody at 4°C overnight. Sections were then washed three times with PBST (0.1% Triton), incubated with secondary antibody (anti-goat, Alexa Fluor 488 Invitrogen cat# A-11055) for 1 hour at room temperature, washed three times with PBST, incubated with Hoechst (1:1000, Thermo Fisher cat# 62249) for 15 minutes at room temperature, and finally washed one more time with PBST before being covered with Fluoromount-G (Southern Biotech cat# 0100-01) and sealed for microscopy.

### Microscopy and Image analysis

OB images were taken using a Leica TCS SP8 confocal microscope equipped with a 20x/.75 objective and Leica LAS X software (RRID:SCR_013673, https://www.leica-microsystems.com/products/microscope-software/details/product/leica-las-x-ls). To obtain the entire volume of each OB section, Z-stacks of ∼30-40um per sample were collected at 1024×1024 resolution with 4x line averaging and 2x frame averaging. All images were stitched within the LAS X software. OB images were analyzed using Imaris software (v10.0, RRID:SCR_007370, http://www.bitplane.com/imaris/imaris). For each image, a region of interest was drawn over the glomerular layer (excluding the olfactory nerve layer) and was used to generate a mask. Volumetric surfaces were then generated following absolute intensity thresholding to account for per-sample background fluorescence signal. To quantify transduction efficiency of each serotype, the total area (um^2^) of TdTomato signal and OMP signal within the mask were quantified. The ratio of TdTomato signal to OMP signal was then calculated. An average of 23.3 OB sections were imaged for each of the 33 mice, for a total of 760 confocal images.

To blind the researchers to which serotype was being analyzed, image files were automatically renamed and randomly sorted into separate folders using a program generated in house. The files were converted back into their identifiable names after all Imaris-based quantification was completed.

For a more comprehensive guide on how to analyze images in this manner, please refer to the following protocols.io DOI: dx.doi.org/10.17504/protocols.io.n2bvjn1rxgk5/v1 To access the complete imaging dataset, please refer to “Comparative Analysis of AAV Serotypes for Transduction of Olfactory Sensory Neurons”, accession number S-BIAD1370 (DOI: 10.6019/S- BIAD1370), on BioImage Archive.

### Tissue harvest and Nuclei Isolation

Four weeks after nasal inoculation with AAV1, AAV7, AAV-DJ/8, and AAVrh10 (mixed together in equal proportions, final concentration of 1.22e11 vg/ml for each serotype), 4 male C57BL/6J mice were deeply anesthetized and transcardially perfused with ice-cold PBS. The OE was then rapidly collected. Immediately following dissection, samples were cut into small pieces and processed using GentleMACS nuclei isolation protocol (Nuclei Extraction Buffer [Miltenyi Biotec, cat# 130-128-024], Protector RNAse Inhibitor [Millipore Sigma, cat# 3335402001], GentleMACS C tubes [Miltenyi Biotec, cat# 130-093-237], GentleMACS Octo Dissociator [Miltenyi Biotec, cat# 130-096-427], MACS SmartStrainers 70um [Miltenyi Biotec, cat# 130-098-462], MACS SmartStrainers 30um [ Miltenyi Biotec, cat# 130-098-458]). In brief, samples were placed in 2mL of Miltenyi Nuclei Isolation Buffer and Protector RNAse Inhibitor in GentleMACS C tubes. Samples then underwent the preprogrammed “nuclei isolation” program in a GentleMACS Octo Dissociator. Immediately after dissociation, samples were strained through a 70um MACS SmartStrainer and collected in a 15ml tube and centrifuged at 500 x g for 5 minutes at 4°C. The supernatant was extracted and discarded, and the resulting pellet resuspended in 1mL of ice-cold PBS. Resuspended samples were then run through a 30um MACS SmartStrainer. Upon visual inspection of nuclei following isolation, it was determined that debris levels were low enough for samples to proceed immediately to library preparation.

### snRNAseq Bioinformatic Analysis

#### Alignment and Quantification of Transcripts

4 sets of paired-end snRNAseq FASTQ files, each corresponding with a technical replicate, were aligned and quantified using STAR v2.7.11b (RRID:SCR_004463, http://code.google.com/p/rna-star/) and STARSolo (RRID:SCR_021542, https://github.com/alexdobin/STAR/blob/master/docs/STARsolo.md)^14^. For STARSolo to generate the spliced/unspliced count matrix, we used the velocyto flag available in the pipeline. All downstream steps were performed using the spliced count matrix unless specified otherwise. To include the AAV serotypes in the alignment, we generated a custom reference FASTA and GTF file using the GENCODE (RRID:SCR_014966, https://www.gencodegenes.org) GRCm39 vM34 release^15^.

#### Ambient RNA Correction

After alignment and quantification, raw spliced, unspliced, and ambiguous count matrices were generated for each sample. To reduce the effects of ambient RNA contamination and filter out empty droplets, we applied CellBender v0.3.0 (https://github.com/broadinstitute/CellBender) to each replicate’s corresponding spliced/mature RNA count matrix^16^. The resulting corrected matrix was transformed into a Scanpy object for further analysis in Python v3.10.14 (RRID:SCR_008394, https://www.python.org).

#### Processing, Quality Control, and Integration of snRNAseq data

Each replicate was processed and filtered according to the standard processing guidelines from Scanpy v1.10.1 (RRID:SCR_018139, https://github.com/theislab/scanpy)^17^. Initial thresholds of minimum 10 cells and 200 genes were set to remove any empty droplets missed by CellBender. Gene/feature initial cutoffs were left more lenient to avoid any possibility of filtering out the AAV genes in case low transduction and expression in the dataset. We set feature/gene and UMI barcode cutoffs to remove dead, low-quality cells, and doublets/multiplets. The cutoffs were set as: 1500 for number of features, 2000 for total counts, and 1% for percent mitochondrial counts. Any cells with gene, count, and mitochondrial count percentages above these thresholds were removed from the dataset. After filtering, samples 1, 2, 3, and 4 had 5810, 6065, 5480, and 5820 cells, respectively.

Doublet removal was performed using the raw count data using Solo from scvi-tools v1.0.4 (https://github.com/scverse/scvi-tools)^18,19^. Filtering, including doublet removal, was performed on each individual sample and after integration of the count matrices. For data integration and batch effect removal, we used the single-cell Variational Inference (scVI) model from scvi-tools. Briefly, the model learns a nonlinear mapping between the latent space and the parameters of a zero-inflated negative binomial distribution used to generate gene expression counts. Batch correction is performed by including batch annotations as an input to the decoder network, allowing the model to learn batch-specific effects that can be removed when sampling from the latent space. Finally, the standard Scanpy processing pipeline including library log-normalization, selection of highly variable genes (N=3500), dimensional reduction with PCA and UMAP, and clustering was applied to the integrated data. We did not use scVI to regress out covariates due to the likelihood of over-correction in the batch-effect removal step. The resulting cell x gene matrix was 20081 x 3500 (24415 raw). When possible, computational speeds were accelerated using an NVIDIA A30 GPU and the rapids single-cell library v0.10.5 (https://rapids-singlecell.readthedocs.io), a library for GPU-acceleration of Scanpy processing steps^20^.

#### Classification of AAV+ Cells

For identification of AAV+ cells, we set a threshold based off the log-normalized expression. Any cell *i* is considered positive for AAV gene *j* if

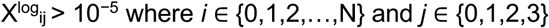

Where *N* is the total number of cells minus one and *j* corresponds to each of the 4 AAV serotypes. By stratifying according to the cell types (Celltype Annotation), we were able to find the cell-specific frequency distribution of all AAV serotypes in the snRNAseq dataset.

#### Cell Type Annotation

Using the latent representation generated by scVI after integration, a uniform manifold approximation projection (UMAP) was generated. Clusters were generated using the Leiden algorithm at multiple resolutions to capture different size populations and cell type specific markers were used to confirm the identities. Using canonical markers derived from the literature, we identified and annotated each cluster according to the cell type that it belonged to (see table below). For clusters that were seemingly distinct but expressed the same cell type clusters, we included a separate metadata column to annotate the clusters with an additional identifier (i.e., mSus and mSus2).

**Table 1.**
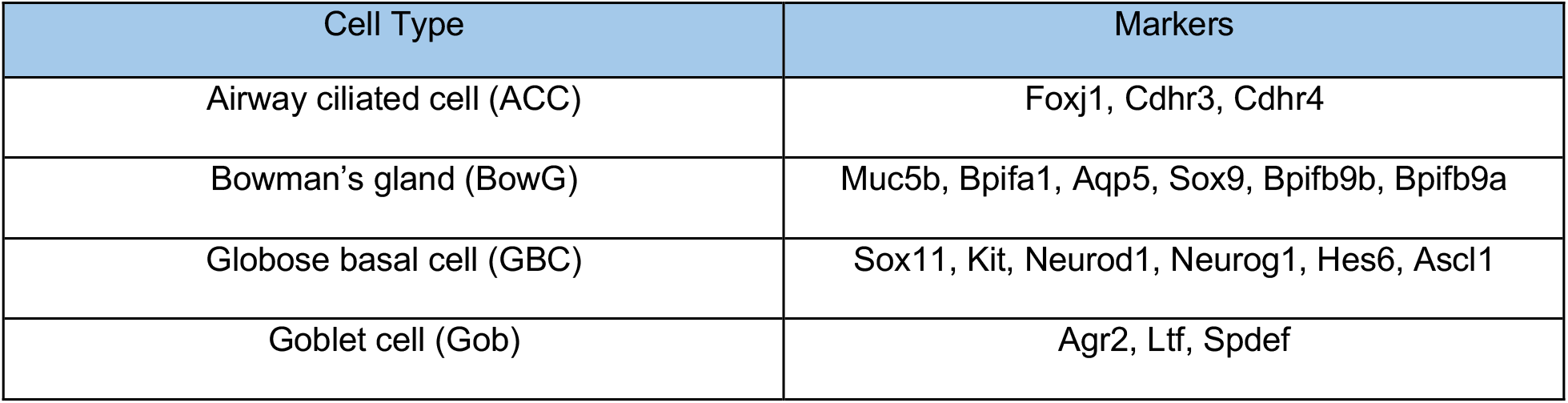
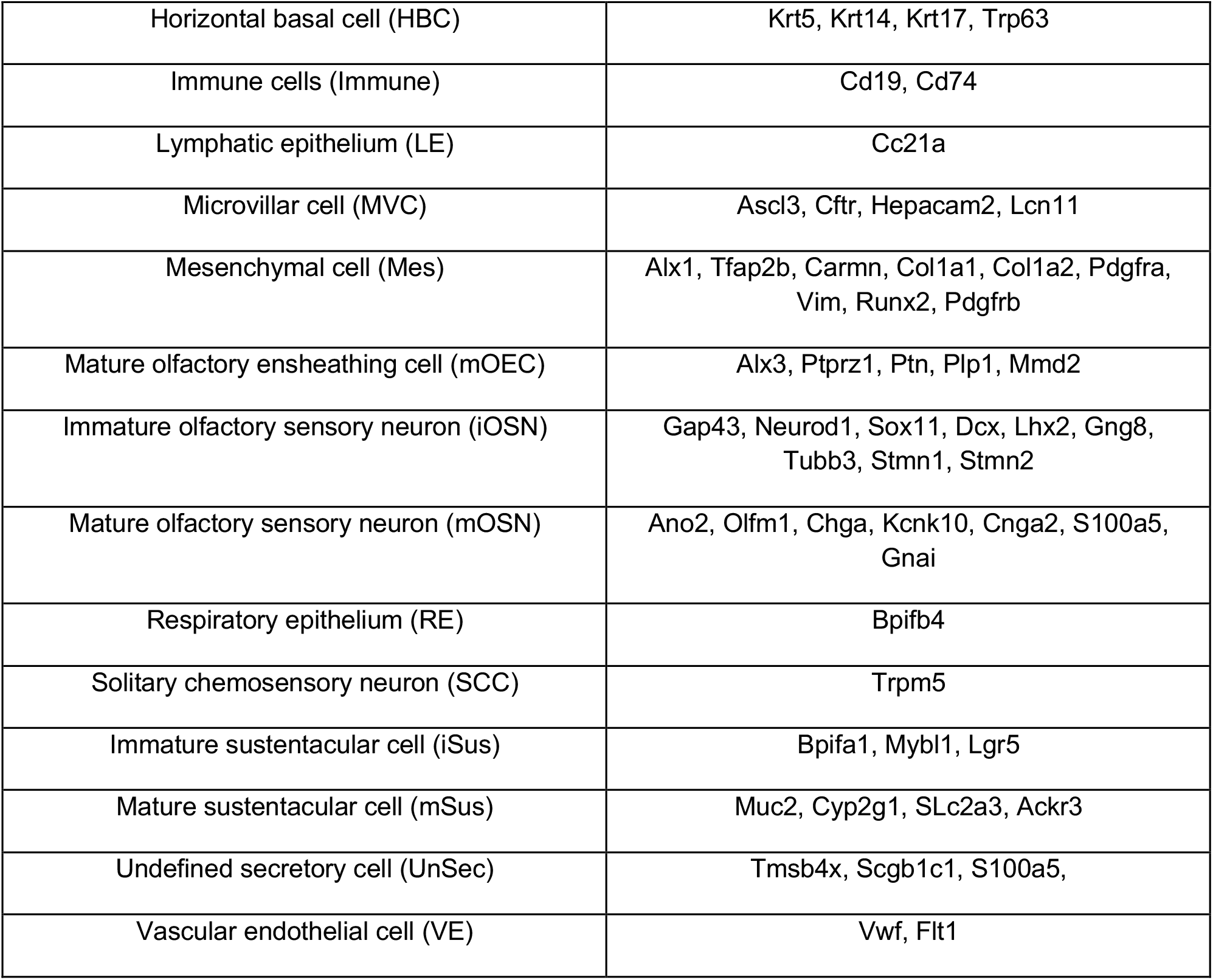
Cell Types and Cell Type Markers.

#### Pseudotime and Terminal Fate Probability

We selected only relevant lineages of interest for downstream analysis and confirmation, specifically OSN, Sus, MVC, and BowG lineages. The relevant cell types include HBC, GBC, iOSN, mOSN, iSus, mSus, MVC, and BowG. After subsetting, the nearest neighbor graph was re-calculated again to find the relationships between the remaining clusters. To improve the visualization of our dataset, we used the minimum distortion embedding (MDE) technique available in pyMDE v0.1.18 (https://pymde.org) to generate a new embedding space^21^.

To confirm the identities of our cell type annotations, we determined the pseudotime relationships between clusters using Palantir^22^ v1.3.3 (https://github.com/dpeerlab/Palantir). We specified HBC as the root cell while defining mOSN, mSus, MVC, and BowG as the terminal points for Palantir. We then examined the Palantir-generated terminal fate probabilities of HBC, GBC, and iSus to confirm the annotated cluster identities. The terminal fate probabilities were evaluated with a 2-sided t-test for each individual lineage and a chi-squared goodness of fit (GOF) to evaluate the total terminal fates with α=0.05.

## Resource Availability

All code used in Bioinformatic Analysis is publicly available on Github at https://github.com/LiuzLab/Mouse-AAV-OSN. Raw count matrices and the final processed AnnData object are available with the Github code on Zenodo (https://doi.org/10.5281/zenodo.13620762). Raw FASTQ files are available on the CRN Cloud.

All images collected for this manuscript are publicly available through BioImage Archive (accession S- BIAD1370, DOI: 10.6019/S-BIAD1370)

CC BY: Comparative Analysis of AAV Serotypes for Transduction of Olfactory Sensory Neurons © 2024 by Benjamin Belfort is licensed under CC BY 4.0. To view a copy of this license, visit https://creativecommons.org/licenses/by/4.0/

For extended materials and methods regarding AAV production and snRNAseq library preparation and sample submission, please refer to the supplemental information.

## Supporting information

Supplemental Information for Belfort et al 2024

## Acknowledgments

This research was funded in whole or in part by Aligning Science Across Parkinson’s [10000400] through the Michael J. Fox Foundation for Parkinson’s Research (MJFF). For the purpose of open access, the author has applied a CC BY public copyright license to all Author Accepted Manuscripts arising from this submission. Special thanks to the Texas Children’s Hospital Neuroconnectivity Core for assistance with viral packaging and Baylor College of Medicine’s Single Cell Genomics Core for assistance with snRNAseq library preparation.

