## Supplemental Information for Belfort et al 2024 for "Comparative Analysis of AAV Serotypes for Transduction of Olfactory Sensory Neurons"

1 Genetics and Genomics Graduate Program, Baylor College of Medicine, Houston, Texas 77030, USA.
2 Department of Molecular and Human Genetics, Baylor College of Medicine, Houston, Texas 77030, USA.
3 Jan and Dan Duncan Neurological Research Institute, Texas Children's Hospital, Houston, Texas 77030, USA
4 Medical Scientist Training Program, Baylor College of Medicine, Houston, Texas 77030, USA.
5 Aligning Science Across Parkinson’s (ASAP) Collaborative Research Network, Chevy Chase, MD 20815.
6 Department of Pediatrics, Section of Neurology, Baylor College of Medicine, Houston, Texas 77030, USA
7 Department of Neurosurgery, Baylor College of Medicine, Houston, Texas 77030, USA
8 Department of Neuroscience, Baylor College of Medicine, Houston, Texas 77030, USA.
*,^ These authors contributed equally

° Corresponding author: Benjamin R. Arenkiel

**Author Contributions:** BDWB and BRA- project conceptualization; JJ and BDWB- bioinformatic analyses; BDWB, AG, AMI- performed research; JPM, JOG- provided essential resources; BDWB, JJ, AG, AMI- performed analyses; BDWB, JJ- wrote the paper; BP, BT, HC- provided feedback and technical support.

**Collaborators:** aSCENT-PD Investigators: Benjamin Arenkiel, Zhandong Liu, Brit Mollenhauer, Josef Penninger, Maxime Rousseaux, Armen Saghatelyan, Natalina Salmaso, Christine Stadelmann, Michael G Schlossmacher, Julianna J Tomlinson, John M Woulfe

**Competing Interest Statement:** Disclose any competing interests here.

**Classification:** Neuroscience

**Keywords:** AAV, gene therapy, olfaction, olfactory sensory neurons

**This PDF file includes:**

Supporting text
Table S1
SI References

**Extended Materials and Methods:**

***AAV Production***
All AAVs were packaged in-house by the Texas Children's Hospital Jan and Dan Duncan Neurological Research Institute's Neuroconnectivity Core, as described below.

Cell Culture and Transfection
A three-vector system (serotype AAV1, AAV2, AAV5, AAV7, AAV8, AAV9, AAV-DJ/8, AAV-PHP.eB, AAV-PHP.S, AAVrh10, AAV-SCH9 was used for AAV production (Cell Biolabs). HEK-293 cells were plated in 15 cm dishes at a density that yielded ~70% confluency the following day. Cells were then transfected in each plate with 25µg helper plasmid, 25µg serotype specific AAV vector, and 25µg of AAV shuttle vector using polyethylenimine (PEI). In a 1:3 ratio (µg DNA:µg PEI), the solution was added dropwise to cells. After 4-6 hours, the medium was changed to DMEM, 5% FBS, 1x Penicillin/Streptomycin. 48-72 hours later, transfected cells were harvested using a cell scraper. Cells were pelleted by centrifugation at 3500rpm for 10 minutes at 4°C. The supernatant was removed, and the pellet resuspended in TMN (50mM Tris pH8.0, 5mM $MgCl_2$, 0.15M NaCl) at a concentration of 1ml/plate. The resuspended cells were frozen at -80°C overnight.

Purification
10ul of DNaseI (10mg/ml) and 10ul RNase A (1mg/ml) were added to each plate of defrosted cells in TMN. Plates were incubated at 37°C for 30 minutes, shaking frequently. 100µl of 5% sodium deoxycholate solution in water was added to each plate and mixed gently. Plates were then incubated at 37°C for 10 minutes. The suspended samples were transferred from the plates to tubes and placed on ice for 15 minutes. The tubes were then centrifuged at 3700rpm for 10 minutes and the supernatant was collected.

Iodixanol Gradient
OptiPrep™ (Millipore Sigma D1556-250ML), or iodixanol, was purchased as a 60% (W/V) stock in water. 15%, 25%, and 40% dilutions of iodixanol were made in PBS-MK (1x PBS, 1 mM $MgCl_2$, 2.5 mM KCl). 2.5µl phenol red solution (0.5% stock in PBS-MK) was added per 1ml of iodixanol solution in the 25% and 60% fractions. The gradient was loaded to the bottom of Beckman OptiSeal 16X67mm tubes (Cat# 362181) starting with 1.5ml 15% iodixanol, 1.3 ml 25%, 1.4ml 40% and finally 1.3ml 60% iodixanol. The supernatant collected from the previous purification step was then placed on top of the gradient. Tubes were centrifuged at 60000rpm for 90 minutes in a Beckman NVT 65 rotor. The clear band below the 60% mark (and below the white cellular debris layer) was collected using a needle and syringe. The collected volume from each tube was approximately 1.5ml.

Concentration
The goal of this step was to remove the OptiPrep and concentrate the AAV using an Amicon Ultra-15 Centrifugal Filter (Millipore Sigma, UFC9 100 24). An Amicon column was equilibrated with 15ml of DPBS (no Mg, no Ca) by centrifugation at 2500rpm for 5 minutes. The collected band from the OptiPrep gradient was mixed with approximately 40ml of DPBS. The samples were run in batches through the Amicon filter, discarding the filtrate between spins. The virus was then washed three times with 15ml of DPBS with centrifugation after each wash at 2500rpm for 10 minutes. The virus was collected to a sterile microcentrifuge tube, aliquoted, and frozen to -80°C.

Viral Titer

Titering of virus was performed using Applied Biological Materials qPCR AAV Titer Kit (Cat# G931) and following the manufacturer's recommended protocol. Viral preparations were first diluted to ~$10^8$ GC/mL before undergoing viral lysis at room temperature for 3 minutes. A standard curve was generated using five 10-fold serial dilutions of provided Standard Control DNA (dilutions 1/100 to 1/100,000). qPCR components and cycling conditions are found within the manufacturer's accompanying product datasheet. Final titer analysis was performed using the manufacturer's provided calculation file.

To view this protocol on protocols.io, please refer to the following DOI:
dx.doi.org/10.17504/protocols.io.81wgbzwj3gpk/v1

***snRNAseq Library Preparation and Sample Submission***

Library Preparation
The single-cell gene expression Library was prepared according to the Chromium Single Cell Gene Expression 3'v3.1 instruction (PN-1000121, PN-1000120, PN-1000213, 10x Genomics). Briefly, single cells, reverse transcription (RT) reagents, Gel Beads containing barcoded oligonucleotides, and oil were loaded on a Chromium controller (10x Genomics) to generate single-cell GEMS (Gel Beads-In-Emulsions) where full-length cDNA was synthesized and barcoded for each single cell. Subsequently, the GEMS were broken and cDNA from each single cell was pooled. Following cleanup using Dynabeads MyOne Silane Beads, cDNA is amplified by PCR. The amplified product was fragmented to optimal size before end-repair, A-tailing, and adaptor ligation. The final library was generated by amplification.

Sequencing
After passing the quality control, the next-generation sequencing of libraries was performed on NovaSeq 6000 (Illumina). All sequencing was performed by Azenta Life Sciences. Below is a table containing the parameters of sample submission.

**Table S1. snRNAseq Submission Parameters**

| | |
|---|---|
| Number of Pools | 4 |
| Total Number of Libraries | 4 |
| Library Size | ~530bp |
| Library Diversity | High diversity |
| Sequencing Configuration | Illumina, 2x150bp, 375Gb |
| Illumina Index Chemistry | Dual index |
| Illumina Index Length | 10bp |
| Sequencing Primers | Standard Illumina |
| Indexing Primers | Standard Illumina |
| Phi-X Spike-In | 5% |
